# Pathogen virulence does not alter antibiotic efficacy under growth-matched *in vivo* conditions

**DOI:** 10.64898/2026.05.19.726418

**Authors:** Victor Wolff Bengtsen, Susanne Häussler, Mohammad Roghanian

## Abstract

Whether pathogen virulence influences antibiotic efficacy independently of bacterial growth dynamics has not been directly tested under controlled *in vivo* conditions. Here, we used the *Galleria mellonella* larval infection model to address this question. We first compared *Pseudomonas aeruginosa, Escherichia coli, Acinetobacter baumannii*, and *Staphylococcus aureus* across a wide range of inoculum sizes, revealing pronounced pathogen-specific differences in virulence. Despite these differences, all pathogens followed a similar infection trajectory characterised by an initial reduction in bacterial burden followed by successful proliferation within the host. Focusing on *P. aeruginosa* and *E. coli*, which differ markedly in virulence but display comparable *in vivo* growth rates (doubling times of approximately 30 minutes), we assessed antibiotic efficacy under experimentally controlled, growth-matched conditions. Treatment with ciprofloxacin or gentamicin at defined multiples of the MIC reduced bacterial loads, prevented host death, and cleared infections to an indistinguishable extent for both pathogens. These findings demonstrate that pathogen virulence does not determine antibiotic efficacy when bacterial growth is comparable *in vivo*, and support the predictive value of MIC-based susceptibility testing during active infection.

## Introduction

Antimicrobial susceptibility testing, most commonly defined by determination of the minimal inhibitory concentration (MIC), is a cornerstone of clinical microbiology and guides antibiotic selection in routine practice [1]. Despite this, antibiotic treatment failure remains a persistent clinical challenge, even when the infecting pathogen is classified as susceptible *in vitro* [2, 3]. Such discordance between laboratory susceptibility results and clinical outcome highlights the complexity of antimicrobial therapy *in vivo* and raises important questions about which infection-related factors influence treatment success.

A fundamental determinant of antibiotic efficacy is bacterial physiology. Actively growing and metabolically active bacteria are generally more susceptible to bactericidal antibiotics, as most antimicrobial agents target processes tightly coupled to biosynthesis and proliferation, including cell wall assembly, DNA replication, transcription, and protein translation [4–6]. In contrast, slow-growing or metabolically restrained cells exhibit reduced killing and may survive exposure through phenotypic tolerance, even in the absence of classical resistance mechanisms [3–10]. Antibiotic tolerance, the ability of genetically susceptible bacteria to survive transient antibiotic exposure due to reduced growth or metabolic activity, is increasingly recognised as a major contributor to treatment failure and a stepping stone toward the evolution of heritable resistance [3, 8–10]. These growth-dependent principles are well established *in vitro* and underpin much of our current understanding of antimicrobial activity.

During infection, however, bacterial growth and metabolism are shaped by host-imposed constraints that are difficult to reproduce under standard laboratory conditions. Nutrient limitation, immune-mediated stress, spatial heterogeneity, and bacterial burden can substantially alter physiological state and thereby influence antibiotic activity [11–13]. For instance, bacteria residing in biofilms or within abscesses experience reduced antibiotic penetration and frequently adopt slow-growing, tolerant phenotypes, while intracellular pathogens may be shielded from extracellular drug concentrations entirely [13–15]. Consequently, it remains unclear whether apparent discrepancies between MIC-based susceptibility testing and treatment outcome reflect intrinsic limitations of susceptibility testing or context-dependent differences in bacterial growth dynamics during infection.

Among the factors proposed to influence treatment failure, pathogen virulence has received increasing attention [16–19]. Highly virulent strains often cause rapid host damage, increased mortality, and severe disease, leading to the assumption that virulence itself may compromise antibiotic efficacy and render highly pathogenic organisms more difficult to eradicate, even when classified as susceptible [20, 21].

Virulence is frequently intertwined with increased bacterial burden, altered metabolic states, and immune evasion, all of which can independently influence antibiotic activity. Indeed, while associations between virulence determinants and antibiotic resistance have been documented in certain pathogen lineages [16–19], a direct causal link between virulence *per se* and reduced antibiotic killing has never been established under controlled *in vivo* conditions in which bacterial growth rates are experimentally matched. Whether virulence alone, independent of its effects on bacterial proliferation and host physiology, alters antibiotic responsiveness therefore remains an open question. Disentangling these variables is essential to determine whether treatment outcome is primarily governed by bacterial growth and metabolic activity, as *in vitro* principles predict, or whether virulence introduces additional complexity that MIC-based testing cannot capture.

Addressing this question requires an *in vivo* model system that allows precise control over infection dose, timing of antimicrobial intervention, and quantitative monitoring of bacterial burden and host survival. The *Galleria mellonella* (greater wax moth) larval infection model has emerged as a versatile platform for studying bacterial pathogenesis and antimicrobial efficacy under physiologically relevant conditions [22–25]. Larvae possess a functional innate immune system with cellular and humoral defence mechanisms that share remarkable similarities with the vertebrate innate immune response [22–25]. Importantly, the model enables systematic comparisons of virulence potential, *in vivo* growth dynamics, and antibiotic treatment outcome within a single, controlled experimental framework [26, 27].

Here, we used the *G. mellonella* model to directly test whether pathogen virulence influences antibiotic efficacy when bacterial growth rates are controlled *in vivo*. We compared clinical strains of *Pseudomonas aeruginosa, Escherichia coli, Acinetobacter baumannii*, and *Staphylococcus aureus* to define pathogen-specific virulence profiles and characterise growth dynamics during infection. Although these pathogens differed markedly in their ability to cause host death, *P. aeruginosa* and *E. coli* exhibited comparable *in vivo* doubling times despite pronounced differences in virulence. Under these growth-matched conditions, treatment with ciprofloxacin or gentamicin at defined multiples of the MIC resulted in equivalent bacterial clearance and host survival, demonstrating that virulence alone does not dictate antibiotic efficacy.

## Results

### Virulence in *Galleria mellonella* is pathogen specific

To establish the *G. mellonella* model as a platform for comparing pathogen virulence under controlled infection conditions, we first examined larval survival following infection with increasing inocula of *P. aeruginosa, E. coli, A. baumannii*, and *S. aureus*. Larvae were infected with defined bacterial doses and monitored for viability over time (Figure 1).

**Figure 1.**
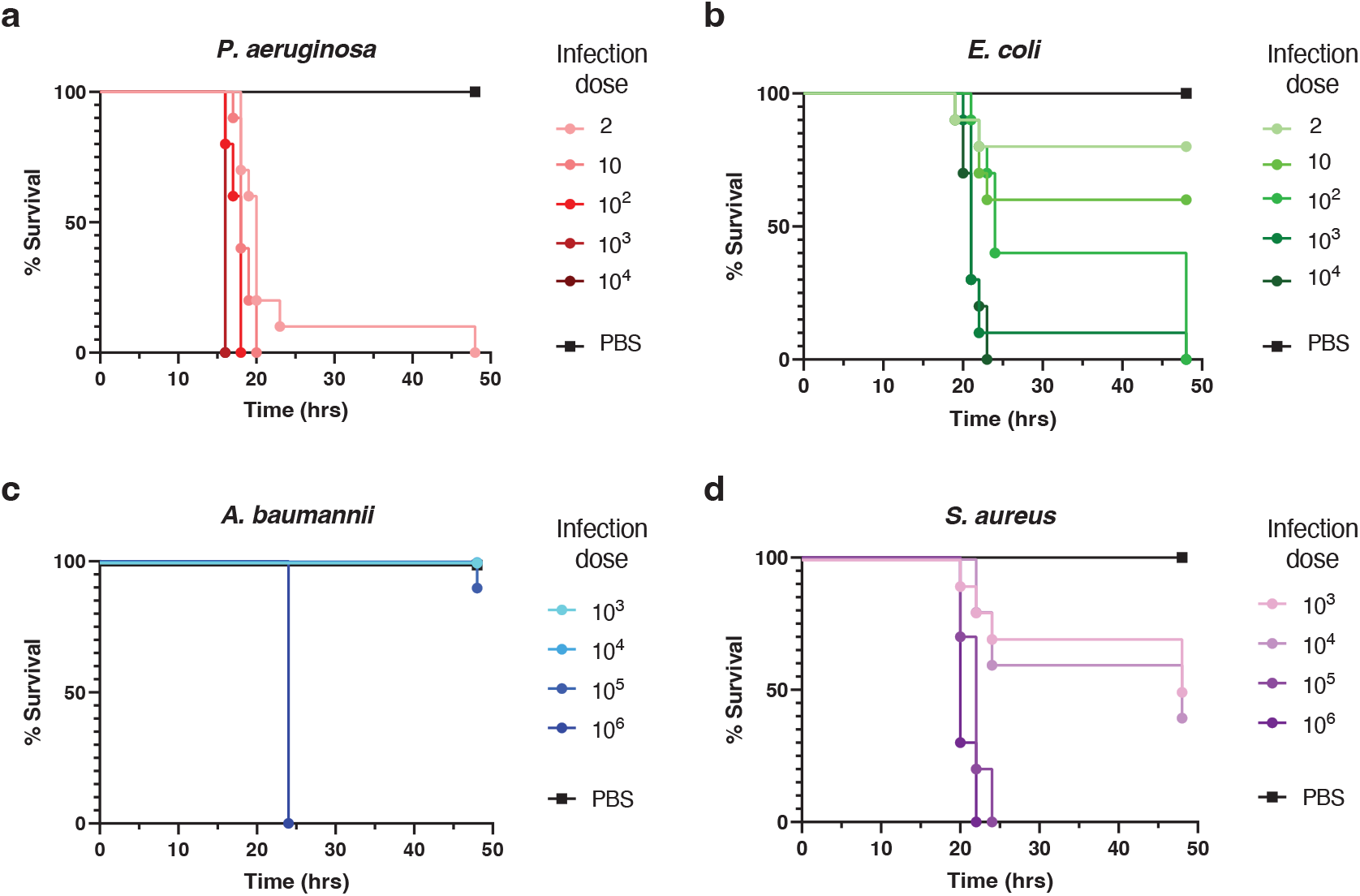
Pathogen-specific virulence profiles in *Galleria mellonella*. Survival of *G. mellonella* larvae following infection with increasing inocula of (a) *P. aeruginosa*, (b) *E. coli*, (c) *A. baumannii*, and (d) *S. aureus*. Data represent mean survival from at least 20 biological replicates per condition. PBS-injected larvae served as controls. Lethal infection requires pathogen-specific inocula that differ by several orders of magnitude.

*P. aeruginosa* exhibited a highly virulent phenotype, with as few as 10 cells consistently causing larval death within 20 hours (Figure 1a). Even lower inocula (2 cells) resulted in complete lethality within 48 hours. In contrast, *E. coli* required substantially higher infection doses to achieve comparable outcomes: 10,000 cells caused death within 24 hours, while doses of ≥100 cells were sufficient to kill all larvae by 48 hours (Figure 1b). Lower inocula (2–10 cells) resulted in partial mortality only.

*A. baumannii* displayed markedly lower virulence in this model. Infection doses of up to 10,000 cells failed to cause larval death, and even 100,000 cells resulted in only limited mortality after 48 hours. Complete lethality was observed only at very high inocula (1,000,000 cells; Figure 1c). Similarly, *S. aureus* required high infection doses to establish lethal infections: inocula of 1,000–10,000 cells caused approximately 50% mortality after 48 hours, whereas ≥100,000 cells resulted in rapid and complete larval death (Figure 1d).

Together, these data demonstrate pronounced, pathogen-specific differences in virulence in *G. mellonella*, with lethal infection requiring inocula that differ by several orders of magnitude across species.

### Establishment and growth of infections in *G. mellonella*

Having identified infection doses that reproducibly caused larval mortality, we next investigated whether the pathogens proliferate within the host and how their *in vivo* growth dynamics compare. Larvae were infected with doses sufficient to cause ≥50% mortality within 24 hours (100 cells for *P. aeruginosa* and *E. coli*; 1,000,000 cells for *A. baumannii* and *S. aureus*), and bacterial loads in the haemolymph were monitored over time (Figure 2).

**Figure 2.**
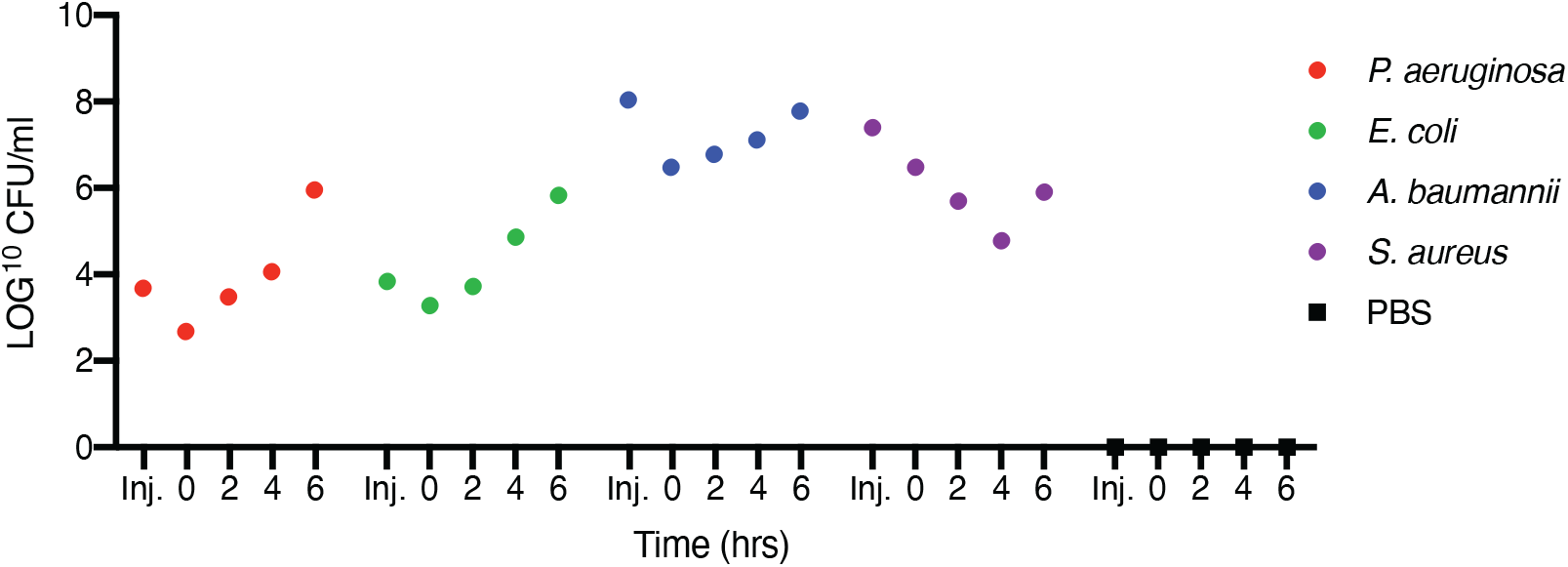
Establishment and *in vivo* growth of bacterial infections in *Galleria mellonella*. Larvae were infected with 100 cells of *P. aeruginosa* or *E. coli*, or 1,000,000 cells of *A. baumannii* or *S. aureus*. Bacterial loads in the haemolymph were quantified at indicated time points. Data represent means from 10 biological replicates. *P. aeruginosa* and *E. coli* exhibit comparable *in vivo* growth rates following a transient clearance phase, indicating similar proliferative and metabolic activity during infection.

Following infection with either *P. aeruginosa* or *E. coli*, we observed an initial reduction in bacterial load within the haemolymph, consistent with early host-mediated clearance. This decrease was transient, and both pathogens subsequently established robust infections characterised by exponential growth (Figure 2). The *in vivo* growth rates of *P. aeruginosa* and *E. coli* were similar, with doubling times of 31.1 ± 3.7 minutes and 26.9 ± 6.6 minutes, respectively. These rates closely matched their growth in rich medium *in vitro* (29.8 ± 0.2 minutes for *P. aeruginosa* and 20.0 ± 0.2 minutes for *E. coli*), indicating comparable proliferative and metabolic activity during infection.

Infections with *A. baumannii* and *S. aureus* displayed a more pronounced initial reduction in bacterial burden, with approximately 2-log and >3-log decreases, respectively, before subsequent recovery and growth (Figure 2). Despite this initial clearance phase, both pathogens were able to adapt to host pressures and establish productive infections.

These findings indicate that successful infection in *G. mellonella* involves an initial adaptation phase followed by bacterial proliferation. Importantly, they reveal that *P. aeruginosa* and *E. coli*, despite marked differences in virulence, exhibit comparable *in vivo* growth rates, providing a controlled physiological context to directly compare antibiotic efficacy independently of growth-related effects.

### Antibiotic treatment effiicacy in *G. mellonella* is independent of pathogen virulence

Given our ability to quantify pathogen growth *in vivo*, we next examined antibiotic efficacy under conditions where bacterial proliferation was comparable. We focused on *P. aeruginosa* and *E. coli*, which differ substantially in virulence but display similar *in vivo* growth rates, allowing assessment of antimicrobial responses independent of growth-related effects on antibiotic killing.

Larvae were infected with 100 cells of either pathogen and allowed to establish infection for one hour before treatment with ciprofloxacin or gentamicin at 10- or 100-fold the minimum inhibitory concentration (MIC; Table 1). Bacterial loads in the haemolymph and larval survival were monitored over time (Figures 3 and 4).

**Table 1.** The minimal inhibitory concentration (MIC) of gentamicin and ciprofloxacin for the *P. aeruginosa* and *E. coli* isolates.

| MIC [μg/ml] | *P. aeruginosa* PA14 | *Escherichia coli* SH_CPH1 |
| --- | --- | --- |
| Gentamicin | 2 | 8 |
| Ciprofloxacin | 0.125 | 0.03125 |

**Figure 3.**
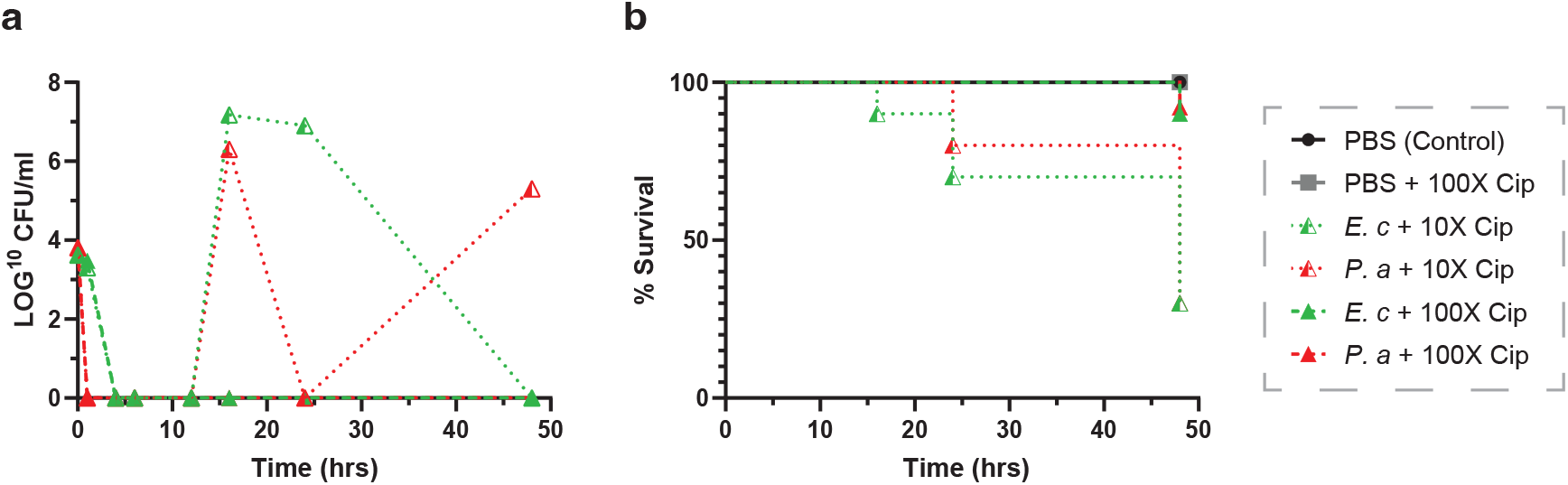
Ciprofloxacin efficacy against growth-matched infections in *G. mellonella*. Larvae infected with *P. aeruginosa* (red) or *E. coli* (green) were treated one-hour post-infection with 10- or 100-fold MIC ciprofloxacin. (a) Bacterial loads in the haemolymph and (b) larval survival were monitored over time. Despite differences in virulence, both pathogens respond similarly to ciprofloxacin when in vivo growth rates are comparable.

**Figure 4.**
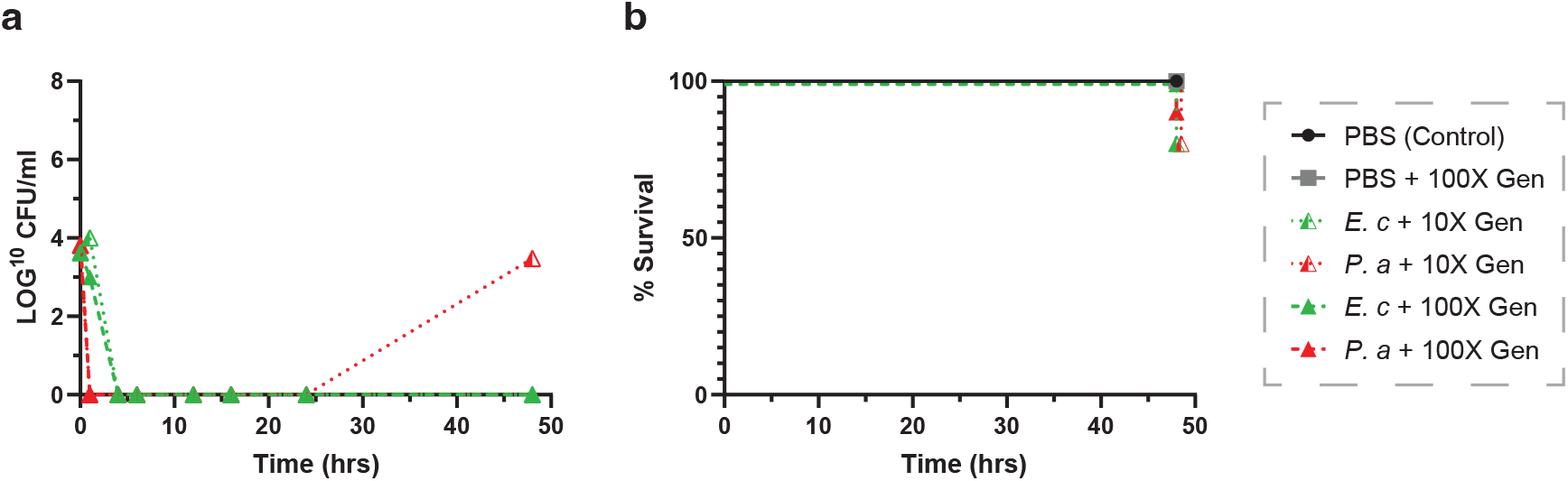
Gentamicin efficacy against growth-matched infections in *G. mellonella*. Larvae infected with *P. aeruginosa* (red) or *E. coli* (green) were treated one-hour post-infection with 10- or 100-fold MIC gentamicin. (a) Bacterial loads in the haemolymph and (b) larval survival were monitored. Gentamicin efficiently clears both pathogens under growth-matched conditions, independent of virulence potential.

Treatment with 10-fold MIC ciprofloxacin significantly reduced bacterial loads for both pathogens but did not fully prevent mortality, resulting in approximately 20% and 70% larval death at 24 and 48 hours post-infection, respectively (Figure 3a–b). In contrast, treatment with 100-fold MIC ciprofloxacin eradicated bacteria from the haemolymph and rescued ~90% of infected larvae for both *P. aeruginosa* and *E. coli* (Figure 3a–b).

Gentamicin was more effective against both pathogens. At 10-fold MIC, gentamicin markedly reduced bacterial burdens and prevented larval death during the first 24 hours, with only ~20% mortality observed by 48 hours (Figure 4a–b). Treatment with 100-fold MIC gentamicin resulted in near-complete clearance of *P. aeruginosa* and complete eradication of *E. coli*, with minimal or no larval mortality (Figure 4a–b).

Injection of 100-fold MIC ciprofloxacin or gentamicin into uninfected larvae had no detectable effect on viability (Figures 3 and 4). Collectively, these data demonstrate that when *in vivo* growth is comparable, despite pronounced differences in virulence, *P. aeruginosa* and *E. coli* respond indistinguishably to antibiotic treatment.

## Discussion

Antibiotic efficacy is often assumed to depend not only on drug properties and bacterial susceptibility, but also on pathogen virulence and infection severity [16–19]. Highly virulent pathogens are frequently thought to be intrinsically harder to eradicate due to enhanced host damage, immune evasion, or altered physiological states during infection [20, 21]. Here, we directly tested this assumption and demonstrate that when bacterial growth is comparable *in vivo*, antibiotic efficacy is independent of pathogen virulence potential.

Using the *G. mellonella* infection model, we observed pronounced differences in virulence across species, with lethal infection requiring inocula that differed by several orders of magnitude. Despite this variation, all pathogens established productive infections characterised by an initial clearance phase, likely reflecting phagocytosis and humoral antimicrobial activity by the larval innate immune system [22–25], followed by exponential proliferation. Crucially, *P. aeruginosa* and *E. coli*, which differ markedly in virulence, exhibited indistinguishable *in vivo* growth rates, providing a controlled physiological framework to assess antibiotic efficacy independently of proliferation-related effects.

Under these conditions, both pathogens responded similarly to ciprofloxacin and gentamicin when administered at equivalent multiples of the MIC. Antibiotic treatment reduced bacterial loads, prevented host death, and cleared infections to comparable extents. These findings indicate that virulence *per se* does not compromise antibiotic-mediated killing once active proliferation is established. Instead, treatment outcome appears to be governed primarily by bacterial growth state and effective drug exposure.

Our results reinforce the well-established principle that antibiotic killing depends on bacterial metabolic activity [4–6]. Ciprofloxacin and gentamicin are bactericidal agents targeting processes tightly coupled to DNA replication and protein synthesis, respectively. Their similar efficacy against growth-matched *P. aeruginosa* and *E. coli* is consistent with the expectation that bactericidal activity scales with active biosynthesis. Conversely, slow-growing or metabolically restrained bacteria are known to survive antibiotic exposure through phenotypic tolerance [3–10], a phenomenon increasingly recognized as a contributor to treatment failure and resistance evolution.

Importantly, our findings bear directly on the predictive value of MIC-based susceptibility testing in clinical practice. The MIC reflects antibiotic activity against exponentially growing planktonic cells under standardised conditions, a physiological state that our growth-matched *in vivo* model closely approximates. Under these conditions, antibiotic efficacy was well predicted by MIC-normalised exposure regardless of virulence, supporting the validity of MIC-guided treatment decisions when bacteria are actively proliferating during infection. Conversely, our data imply that when MIC-based predictions fail clinically, the most likely explanations are pharmacokinetic limitations, impaired drug penetration, spatial heterogeneity, biofilm formation, or transitions to non-replicating states — rather than virulence *per se*. This distinction has practical relevance: it suggests that efforts to improve treatment outcomes in infections caused by highly virulent pathogens should focus on optimising drug exposure and overcoming growth-state-dependent tolerance, rather than assuming that virulence itself renders standard susceptibility thresholds uninformative. In this sense, our findings argue against the clinical intuition that a highly virulent but susceptible pathogen warrants empirical dose escalation beyond MIC-guided recommendations, at least under conditions of active bacterial replication.

Several limitations should be acknowledged. The *G. mellonella* model relies exclusively on innate immunity and does not capture adaptive immune responses that may interact with virulence factors in mammalian hosts. Our antibiotic comparisons were restricted to two Gram-negative pathogens and two bactericidal agents whose activity depends on active growth; whether similar independence of virulence holds for bacteriostatic agents or in chronic biofilm-associated infections remains to be determined. In addition, our analyses were performed at the population level and do not resolve single-cell heterogeneity that may contribute to persistence during therapy.

In summary, our study demonstrates that pathogen virulence does not intrinsically diminish antibiotic efficacy when bacterial growth is controlled *in vivo*. Our findings support a model in which treatment outcome is primarily determined by the intersection of bacterial physiology and drug exposure relative to susceptibility thresholds, rather than by virulence *per se*.

## Materials and Methods

### Bacterial strains

The bacterial isolates used in this study are listed in Table 2. *P. aeruginosa* PA14 is a well-characterised, highly virulent laboratory reference strain; *E. coli* SH_CPH1 is a clinical isolate from the laboratory collection; *A. baumannii* SSI1104 was previously described [28]; and *S. aureus* USA300 is a community-associated methicillin-resistant *S. aureus* reference strain [29]. All strains were routinely cultured in Luria-Bertani (LB) broth (per litre: 10 g tryptone, 5 g yeast extract, 7.5 g NaCl) at 37°C with shaking (180–200 rpm) unless otherwise stated.

**Table 2.** Bacterial isolates used in this study.

| Species | Strain | Source |
| --- | --- | --- |
| *Escherichia coli* | SH_CPH1 | Lab Collection |
| *Pseudomonas aeruginosa* | PA14 | Lab Collection |
| *Acinetobacter baumannii* | SSI1104 | Lab Collection |
| *Staphylococcus aureus* | USA300 | Lab Collection |

### *G. mellonella* infection model

*G. mellonella* larvae were obtained from a commercial supplier (Zoo Center, Copenhagen, Denmark). Healthy larvae weighing approximately 0.25 g and lacking visible signs of melanisation were selected for experiments.

Infection assays were performed as previously describe [30]. In brief, bacterial inocula were prepared from overnight cultures grown in LB broth. Cultures were washed once in phosphate-buffered saline (PBS) and adjusted to the required cell densities by serial dilution. Inoculum accuracy was verified by plating serial dilutions on LB agar. Each larva was injected with 20 µL of bacterial suspension into the last left proleg using a 500 µL Hamilton syringe fitted with a 30-gauge needle. PBS-injected larvae served as negative controls.

Following infection, larvae were incubated at 37°C in the dark. Larval death was defined as melanisation and absence of movement in response to tactile stimulation. Survival was monitored at the indicated time points.

### Growth measurements

To assess bacterial growth during infection, larvae were infected with 100 cells of *P. aeruginosa* or *E. coli*, or with 1×10^6^ cells of *A. baumannii* or *S. aureus*. Infected larvae were incubated at 37°C. At the indicated time points, ten viable larvae were randomly selected and punctured with a sterile needle in the last left proleg. Haemolymph was collected by gentle pressure, pooled, and mixed by gentle pipetting. Bacterial loads were determined by plating tenfold serial dilutions on blood agar plates, incubating overnight at 37°C, and counting colony-forming units (CFU).

*In vivo* doubling times were calculated from the exponential growth phase by fitting a linear regression to semi-logarithmic plots of CFU versus time. Growth rates (k) were derived from the slope, and doubling times were calculated as td = ln(2)/k. *In vitro* doubling times were determined from growth curves in LB broth at 37°C under shaking conditions.

### Antibiotic treatment *in vivo*

Antibiotic treatment was initiated one-hour post-infection. To facilitate handling, larvae were cooled at −20°C for 15 minutes followed by 4°C for an additional 15 minutes. Based on previously reported haemolymph volumes per larva [26], antibiotics were administered in a 10 µL injection volume to achieve final concentrations corresponding to 10-fold or 100-fold the MIC. Antibiotics were injected into the last right proleg using a Hamilton syringe. PBS was injected as control.

At each indicated time point following treatment, haemolymph was collected from six viable larvae and bacterial loads were quantified as described above. Larval survival was monitored in parallel.

### MIC determination

The minimum inhibitory concentrations (MICs) of ciprofloxacin and gentamicin for *P. aeruginosa* PA14 and *E. coli* SH_CPH1 were determined in triplicate using broth microdilution in accordance with EUCAST guidelines, with the following modifications: Overnight cultures grown in LB broth were diluted to an optical density at 600 nm (OD_600_) of 0.005 and exposed to twofold serial dilutions of the antibiotics in microtiter plates. Plates were incubated statically at 37°C for 18–20 hours, and growth inhibition was assessed visually. The MIC was defined as the lowest concentration that completely inhibited visible growth.

## Author contributions

M.R. and S.H. coordinated the study and drafted the manuscript. M.R. and S.H. designed experiments and M.R. analysed the data. M.R. and V.W.B. performed the experiments. All authors agree to be accountable for all aspects of the work.

## Data availability

All data generated or analysed during this study are included in this published article. Strains are available on request.

